# Salivary sex hormone levels following oxytocin administration in autistic and typical women

**DOI:** 10.1101/679282

**Authors:** Tanya L. Procyshyn, Michael V. Lombardo, Meng-Chuan Lai, Bonnie Auyeung, Sarah K. Crockford, Julia Deakin, Sentil Soubramanian, Akeem Sule, Simon Baron-Cohen, Richard A. I. Bethlehem

## Abstract

Oxytocin administration, which may be of therapeutic value for social disabilities, likely influences endogenous levels of other socially-relevant hormones. However, to date, the effects of oxytocin administration on endogenous hormones have only been examined in typical males. The need to consider multi-hormone interactions is particularly warranted in oxytocin trials for autism due to evidence of irregularities in both oxytocin and sex steroid systems. Here, as part of a larger trial with a double-blind cross-over design, we assessed salivary testosterone and oestradiol levels in 16 autistic and 29 typical women before and after intranasal administration of 24IU oxytocin or placebo. Distinct patterns of change in testosterone and oestradiol across time were observed between groups, with autistic women showing increases in both hormones 90 min post-administration and typical women showing small decreases (mean %change oestradiol: +12% Autism, −10% Typical, 95%CI of difference: 5.0–39.4%, p=0.01; mean %change testosterone: +8% Autism, −14% Typical, 95%CI of difference: 7.8–35.6%, p=0.002). Under the oxytocin condition, the group difference in %change testosterone was amplified (+14.4% Autism, −15.2% Typical, p=0.018). Although baseline hormone levels did not differ between groups, greater baseline oestradiol relative to testosterone was negatively correlated with autistic-like traits (r= −0.36, p=0.019) and positively correlated (r=0.35, p=0.02) with self-reported empathy in the overall sample. These results provide further evidence that oxytocin influences endogenous testosterone, with autistic women showing increases similar to previous reports in typical men. These findings may help to identify autistic people expected to benefit most from interventions involving oxytocin.

## Introduction

The neuropeptide hormone oxytocin is known to modulate social behaviour across mammals including humans [1]. For this reason, oxytocin has attracted interest for its potential therapeutic applications in psychiatric conditions characterized by difficulties in social behaviour [2,3]. Intranasal administration, a means of drug delivery to the brain [4], has become the standard method of assessing the effect of a single hormone such as oxytocin on behaviour. Despite recognition of the complexities of the neuroendocrinology underlying human social behaviour and cognition [5], oxytocin administration studies rarely consider the potential influence of other socially-relevant hormones when interpreting the results.

Short-term manipulation of oxytocin is likely to influence endogenous release of other hormones expected to exert their own effects on social behaviour [6]. For example, increased testosterone levels have been reported in men who received oxytocin nasal spray versus placebo [7,8], and levels of arginine vasopressin, a neurohormone closely related to oxytocin, increased in men and women following oxytocin administration [9]. Furthermore, the effects of oxytocin administration on parenting-related behaviors have been reported to depend on baseline endogenous testosterone levels in both sexes [7,10]. Interestingly, oxytocin and testosterone administration are noted to have opposing effects on various social behaviours in typical populations and to show opposite patterns of alteration in psychiatric conditions such as autism and schizophrenia [11], although both hormones have rarely been assessed within the same individuals. While such multi-hormone interplay and its relevance to human social behaviour are yet to be fully elucidated, interactions among oxytocin and sex steroid hormones are well documented in animal research.

Using *in vitro* receptor autoradiography, testosterone has been shown to inhibit oxytocin receptor binding in the brains of male mice [12]. Castration increases the number of oxytocin immunoreactive neurons in the paraventricular nucleus of male mice, while castration plus testosterone implants decreases this number [13]. In rats, Leydig cells cultured with oxytocin or an oxytocin agonist produce higher levels of testosterone, and this increase in testosterone production is mediated by the oxytocin receptor [14]. Oestradiol treatment in ovariectomised rats alters the distribution of oxytocin immune-stained neurons and oxytocin levels in brain regions including the lateral septum, striatum, and amygdala [15]. Pre-treatment with oestradiol enhanced the anxiolytic effect of oxytocin administration in female mice, possibly via enhancement of oxytocin binding density [16]. The likelihood of similar interactions between oxytocin and steroid hormones in humans is supported by an *in vitro* study of neuroblastoma cells demonstrating that the androgen receptor mediates down-regulation of oxytocin gene expression [17]. Taken together, these studies suggest a broadly inhibitory relationship between testosterone and oxytocin and a broadly synergistic relationship between estrogens and oxytocin, although these relationships may be further complicated by sex differences in steroid hormone levels and oxytocin receptor systems in the brain [18].

People with autism spectrum conditions (henceforth autism) are an important clinical group to inform the exploration of the interplay among hormones and its effects on social behaviour. The social and communication challenges that characterize autism [19] have been associated with lower endogenous oxytocin levels in autistic children [20–22]. Furthermore, several lines of evidence support that elevation of sex steroid hormones like testosterone increases the likelihood of autism [23–25]. Nevertheless, sex steroid hormones have not been considered in previous randomized controlled trials assessing the effects of oxytocin administration on social cognition in autistic individuals [26].

To date, the effects of oxytocin administration on endogenous steroid hormone levels have only been examined in typical males. Weisman et al. [7] reported alterations in fathers’ salivary testosterone levels after oxytocin administration relative to placebo. Gossen et al. [8] reported alterations in serum testosterone and progesterone, but not oestradiol, in typical men after oxytocin administration. Increases in testosterone levels after central oxytocin administration have also been reported in male squirrel monkeys [27]. Whether women show similar changes in steroid hormone levels following oxytocin administration, and if the same changes occur in individuals with an autism diagnosis, has not been examined. Given sex differences in the effects of oxytocin on neural activity and social behaviour [27,28], sex differences in the patterns of change in sex steroid hormones after oxytocin administration require clarification. To better assess oxytocin’s potential as a therapeutic agent to improve social functioning in autism, further study of the interplay among oxytocin, other socially relevant hormones, and biological sex is warranted.

To address these questions, we analysed salivary oestradiol and testosterone levels in autistic and typical women before and after intranasal administration of oxytocin or placebo. The primary aim of this study was to assess changes in endogenous sex steroids after oxytocin administration and whether these changes differed with autism diagnosis. Additionally, these data were used to compare baseline sex steroid levels between autistic and typical women and to evaluate relationships of these hormones with two psychological variables linked to autism and sex steroid levels, namely the Autism-Spectrum Quotient (AQ) [30] and Empathy Quotient (EQ) [31]. Although there is no *in vivo* work on which to base predictions for women, given the animal literature and possibility of hypermasculinized phenotype in autism [32,33], we predict that oxytocin administration will promote testosterone decreases in typical women and testosterone increases in autistic women—similar to previous reports for men. Further, we predict a higher balance of testosterone relative to oestradiol (E2:T ratio) in autistic women and/or women with higher levels of autistic-like traits.

## Methods

### Participants

A total of 45 women aged 18–50 years participated in this study. Of these, 16 had a diagnosis of autistic disorder/childhood autism or Asperger’s disorder/syndrome based on DSM-IV or ICD-10 criteria (Autism group) and 29 were typical (Typical group). The two groups did not differ significantly in age or Full IQ (Table 1). No participant had a history of psychotic disorders or substance use disorder, any genetic syndrome associated with autism, intellectual disability, pregnancy, epilepsy, hyperkinetic disorder, Tourette’s syndrome, or current or past use of anti-psychotic, glucocorticoid, psychostimulant, or antihypertensive drugs. Use of hormonal contraceptives and anti-depressants was permitted, as a significant proportion of the study population was expected to be taking such medication.

**Table 1.** Demographic characteristics, psychological questionnaire scores, and baseline hormone levels in the Autism and Typical groups. Values are mean ± SD, unless otherwise specified.

|  | **Autism** | **Typical** | ***p-value*** | ***Cohen's d*** |
| --- | --- | --- | --- | --- |
| **n** | 16 | 29 |  |  |
| **Demographics** |  |  |  |  |
| Age (years) | 29.9 ± 8.4 | 27.2 ± 8.1 | 0.31 | 0.33 |
| Full-IQ<sup>1</sup> | 126.4 ± 20.0 | 113.3 ± 13.7 | 0.14 | 0.87 |
| Hormonal contraceptive use (n) | 0 | 8 |  |  |
| **Psychological variables** |  |  |  |  |
| Autism-Spectrum Quotient (AQ) | 37.1 ± 5.1 | 14.1 ± 7.5 | < 0.01 ** | 3.4 |
| Empathy Quotient (EQ) | 20.7 ± 11.7 | 54.6 ± 14.0 | < 0.01 ** | -2.5 |
| **Baseline hormone levels<sup>2</sup>** |  |  |  |  |
| Baseline oestradiol (pg/ml) | 1.0 ± 0.3 | 1.2 ± 0.5 | 0.055 | -.54 |
| Baseline testosterone (pg/ml) | 70.3 ± 24.9 | 66.4 ± 22.0 | 0.60 | 0.17 |

<sup>1</sup> Wechsler Abbreviated Scale of Intelligence
<sup>2</sup> Baseline hormone levels were calculated as the mean of the two pre-administration samples per participant.
\*\* $p < 0.01$

### Experimental design

This study was approved by the NHS Research Ethics Service (NRES Committee East of England-Cambridge Central, reference 14/EE/0202). All participants provided written informed consent prior to experiments.

Data were collected as part of a larger neuroimaging study [34] with a double-blind, placebo-controlled, within-subjects crossover design. Each participant was randomly assigned to receive placebo or oxytocin first, with the second session taking place a minimum of one week later. For normally cycling women, both sessions were scheduled during the luteal phase of the menstrual cycle.

On the day of the experiment, participants underwent a short health screening by a trained clinician and were deemed fit to participate. Participants received either a dose of 24 IU oxytocin (Syntocinon, Novartis, Switzerland) or placebo (prepared by Newcastle Specials Pharmacy Production Unit) and were instructed to self-administer three puffs to each nostril. Following administration, participants rested for approximately 20 minutes.

Psychological self-report questionnaires were completed in advance of the experiment day. The Autism-Spectrum Quotient (AQ) assesses autistic-like traits in adults of normal intelligence [30], while the Empathy Quotient assesses cognitive and affective domains of empathy [35].

### Saliva collection

Participants were instructed to refrain from consuming caffeinated beverages the day of the experiment and to refrain from consuming alcohol for 24 hours prior to testing. Saliva samples were collected at three timepoints: (1) prior to administration (baseline), (2) shortly after intranasal administration of placebo or oxytocin (6 ± 4 min post-administration), and after completion of experimental fMRI tasks not reported here (96 ± 19 min post-administration). In total, six saliva samples (three under the oxytocin condition, three under the placebo condition) were collected per participant. Saliva samples were collected by passive drool and frozen immediately at −80 °C until analysis.

### Salivary hormone analysis

Salivary hormone level analysis is a minimally invasive method commonly used in behavioural research [36]. Salivary oestradiol and testosterone were analysed using commercially-available enzyme-linked immunosorbent assay (ELISA) kits designed specifically for use with saliva (Salimetrics, USA). Assays were performed by the NIHR Cambridge Biomedical Research Centre, Core Biochemical Assay Laboratory. Each saliva sample was analyzed in duplicate. Most samples showed high internal reliability (average coefficient of variation (CV) < 10% for testosterone and < 15% for oestradiol); samples with a higher CV were successfully re-analysed. The inter-assay CVs for the low and high control were 9% and 7% for the oestradiol assays, respectively, and 12% and 10% for the testosterone assays.

For two samples, the obtained oestradiol concentration was below the detection limit of the kit (<0.1 pg/ml); as oestradiol values for other samples from those individuals were well above the detection limit, the low values were deemed invalid measurements and excluded from analysis. One baseline testosterone value and one baseline oestradiol value were identified as outliers (±3 standard deviation (SD) of the mean). As these measurements appeared to be valid (i.e., other samples from the same participant were at the higher range), they were set as the next highest value that was not an outlier.

Baseline hormone levels were calculated as the mean of the two samples collected before administration per participant. To assess the balance of the two hormones, in line with best practices [37], the baseline oestradiol to baseline testosterone ratio (E2:T ratio) was computed as log(baseline oestradiol) – log(baseline testosterone). To explore changes in hormone levels across time, the percent change relative to baseline ((final - initial)/ initial) was calculated for time point 2 (~5 min post-administration) and time point 3 (~90 min post-administration).

### Statistical analysis

Data are presented as mean and standard deviation. Baseline hormone levels and psychological variables were compared between the Autism and Typical groups using Welch’s t-test. First, paired t-tests were used to assess changes in hormone levels within participants over time. Pre-to post-administration changes in salivary oestradiol and testosterone levels between groups (Autism or Typical) and drug condition (oxytocin or placebo) were then assessed by analysis of variance (ANOVA). Tukey’s honest significant difference (HSD) test was used for post-hoc tests, which maintains the familywise error at p = 0.05 for multiple comparisons. Because linear regression can be sensitive to small datasets with high variability, robust regression using iterated re-weighted least squares was also performed. Robust regression attempts to ignore or down-weight unusual data [38], offering further support that results are not driven by a small number of highly influential datapoints. Statistical analyses were performed using R version 3.5.1 [39]. Effect sizes were calculated using the “effsize” package and the “MASS” package was used for robust regression.

## Results

### Participant characteristics and baseline hormone levels

A comparison of demographic characteristics, questionnaire scores, and baseline hormone levels between the Autism and Typical groups is presented in Table 1. The two groups did not differ significantly in terms of age, IQ, baseline oestradiol, or baseline testosterone. Psychological variables differed between groups, with autistic women having substantially higher AQ scores and lower EQ scores than typical women. Baseline oestradiol and testosterone levels were not significantly related to age (r < 0.10, p > 0.45 for both) or hormonal contraceptive use (Welch’s t-test, p > 0.14 for both hormones). Exclusion of the eight typical women who reported taking hormonal contraceptives did not significantly change the results of the group comparison (Supplementary Table S1).

Next, we explored the relationship between balance of baseline hormones (E2:T ratio) and psychological traits. As shown in Figure 1, greater oestradiol relative to testosterone was negatively correlated with AQ score (r = −0.35, p = 0.018) and positively correlated with EQ score (r = 0.35, p = 0.02) in the overall sample. Similar patterns were present when correlations were determined separately for the two groups (Autism; r = −0.22, p = 0.43 for AQ, r = 0.45, p = 0.10 for EQ; Typical: r = −0.25, p = 0.24 for AQ, r = 0.14, p = 0.43 for EQ), although statistical significance was not achieved with the smaller sample sizes.

**Figure 1.**
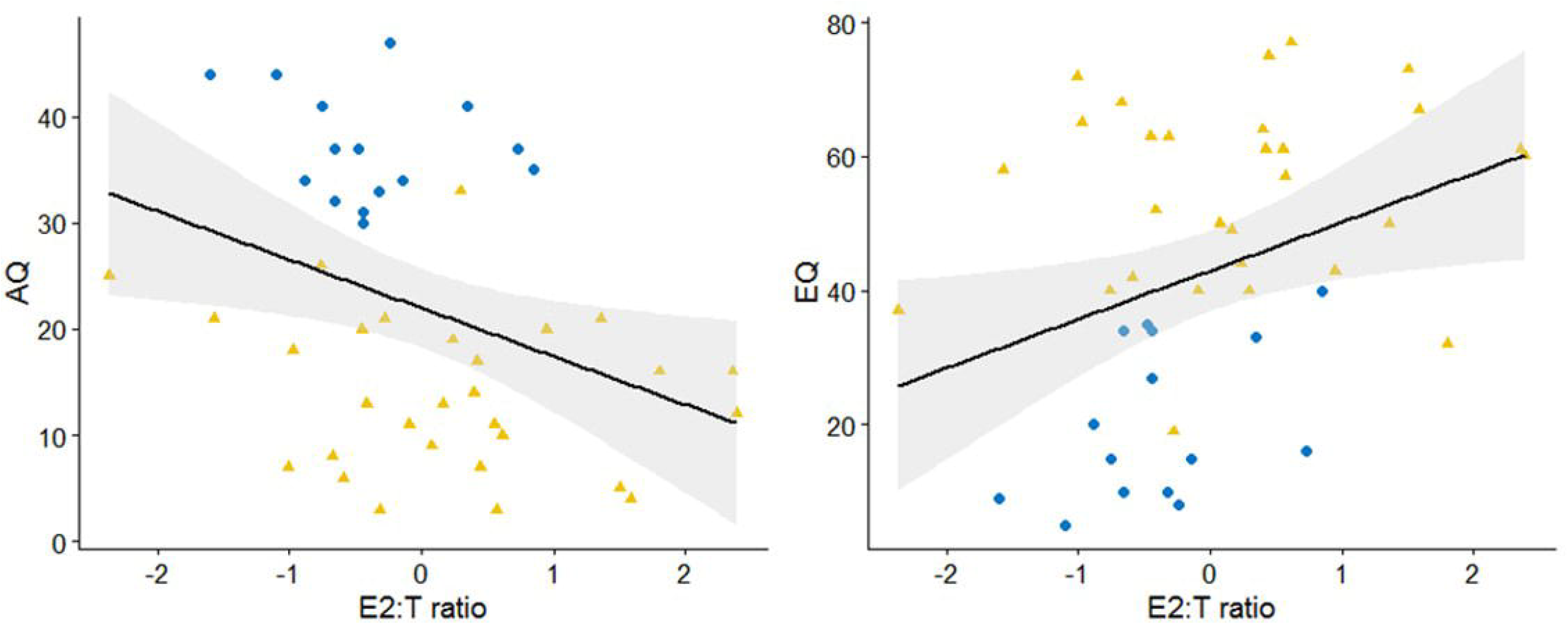
Correlation between balance of baseline oestradiol to testosterone (E2:T ratio) and AQ (left panel) and EQ (right panel). Regression line is indicated in black with 95% confidence intervals shaded in grey. Autistic and Typical women participants are indicated by the darker circles and lighter triangles, respectively.

### Pre-to post-administration changes in salivary oestradiol and testosterone

Mean salivary oestradiol and testosterone levels across the three time points, separated by group and drug condition, are presented in Figure 2. Changes in oestradiol and testosterone levels for individual participants across the three time points are shown in Supplementary Figures S1 and S2, respectively. For the overall sample, both oestradiol and testosterone showed small but significant decreases over the approximately 90 minutes of the experiment (paired t-tests for all participants (n=45), oestradiol: time 1 vs time 2, mean change of - 0.16 pg/ml, *p* <0.001, time 1 vs time 3, mean change of – 0.14 pg/ml, *p*=0.01; testosterone: time 1 vs time 2, mean change of - 10.1 pg/ml, *p* <0.001, time 1 vs time 3, mean change of −7.8 pg/ml, *p* <0.01). However, as shown by Figure 2, the mean hormone levels over time appear to differ between groups and drug conditions.

**Figure 2.**
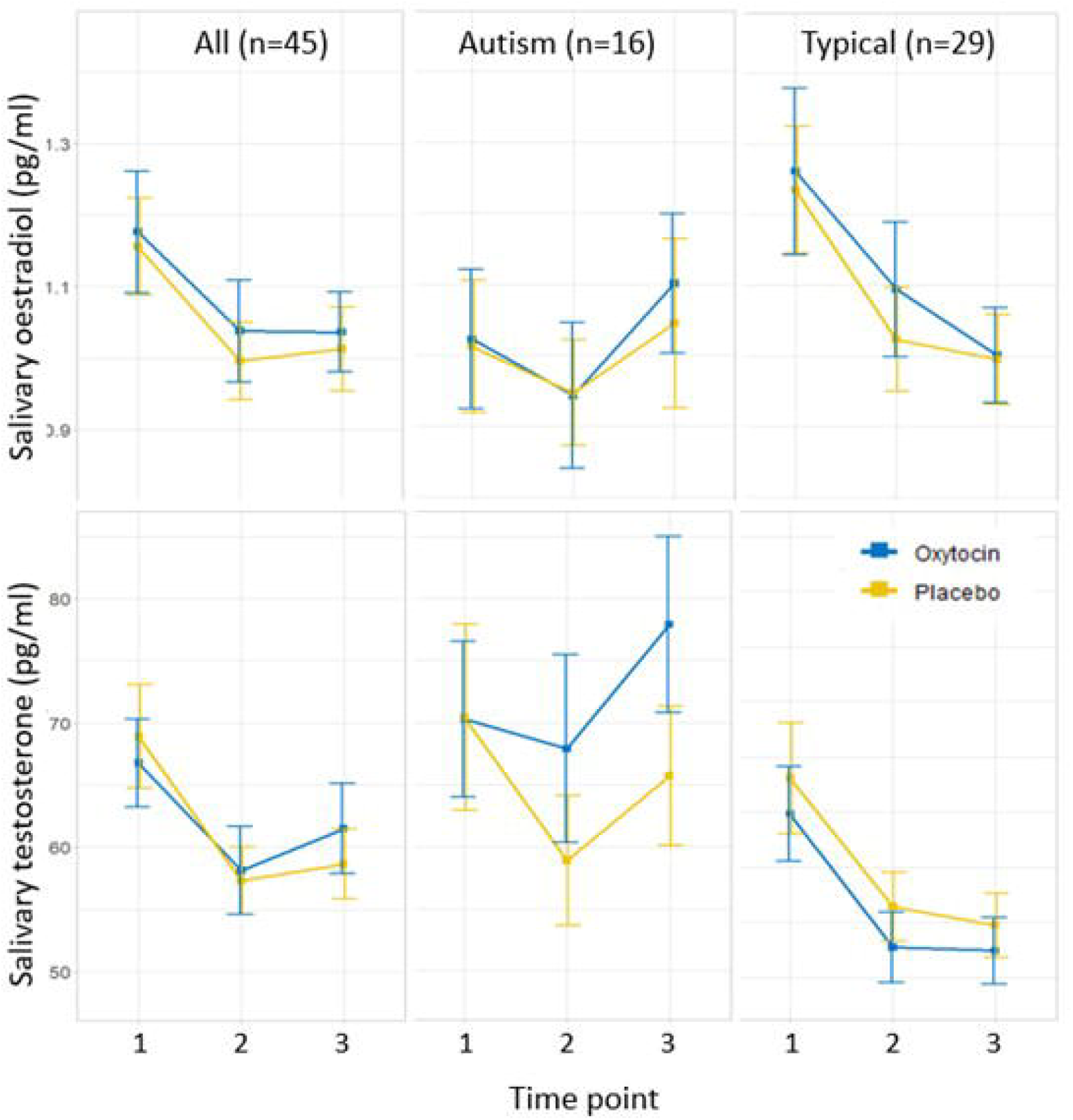
Mean salivary oestradiol levels (upper panels) and testosterone (lower panels) under the placebo and oxytocin conditions across the three measurement points (Time point 1 = before administration, 2= ~5 min post-administration, 3 = ~90 min post-administration). Error bars indicate the standard error of the mean. From left to right, the panels show: (i) hormone levels separated by drug condition for all participants; (ii) hormone levels separated by drug condition for the Autism group; and (iii) hormone levels separated by drug condition for the Typical group.

To assess the effects of group (Autism or Typical) and drug condition (oxytocin or placebo), as well as their interaction, on pre-to post-administration oestradiol and testosterone levels, 2×2 ANOVA was performed. Given the individual variation in hormone levels, the percentage change from baseline was used in the analyses to normalize the data at an individual level.

From time point 1 to 2 (~5 min after administration), there were no significant effects of drug condition or group on either oestradiol or testosterone levels (Supplementary Table 1 and 2, respectively). From time point 1 to 3 (~90 min after administration), a significant group effect, but not a drug condition effect or group × drug interaction, was found for %change oestradiol (Table 2). The mean %change oestradiol from time point 1 to 3, not accounting for drug condition, was +12% for the Autism group and −10% for the Typical group (Tukey HSD, p = 0.01, 95% confidence interval of difference: 5.0–39.4%). A significant group difference was also found for %change testosterone from time point 1 to 3 (Table 2). The mean %change in testosterone, not accounting for drug condition, was +7.8% for the Autism group and −13.9% for the Typical group (Tukey HSD, p = 0.002, adjusted for multiple comparisons, 95% confidence interval of difference: 7.8–35.6%). Under the oxytocin condition the difference in %change testosterone from time point 1 to 3 between groups was even larger (Figure 3), with a mean change of +14.4% for the Autism group and −15.2% for the Typical group (Tukey HSD, p = 0.018, 95% confidence interval of difference: 3.8–55.6%).

**Table 2.** ANOVA tables of changes in salivary oestradiol and testosterone from time point 1 (before administration) to time point 3 (~90 min post-administration).

|  | % Change Oestradiol |  |  |  |  | % Change Testosterone |  |  |  |  |
| --- | --- | --- | --- | --- | --- | --- | --- | --- | --- | --- |
|  | Sum Sq<sup>1</sup> | df | F | p | eta-squared | Sum Sq<sup>1</sup> | df | F | p | eta-squared |
| **Drug condition** | 0.039 | 1 | 0.25 | 0.62 | 0.071 | 0.02 | 1 | 0.20 | 0.65 | 0.002 |
| **Group** | 1.02 | 1 | 6.63 | 0.012* | 0.003 | 0.97 | 1 | 9.66 | 0.003** | 0.10 |
| **Group × Drug** | 0.011 | 1 | 0.07 | 0.80 | 0.001 | 0.13 | 1 | 1.30 | 0.26 | 0.01 |
| **Residual** | 13.2 | 86 |  |  |  | 8.7 | 86 |  |  |  |

<sup>1</sup> Type II sum of squares.
\* $p < 0.05$ , \*\* $p < 0.01$

**Figure 3.**
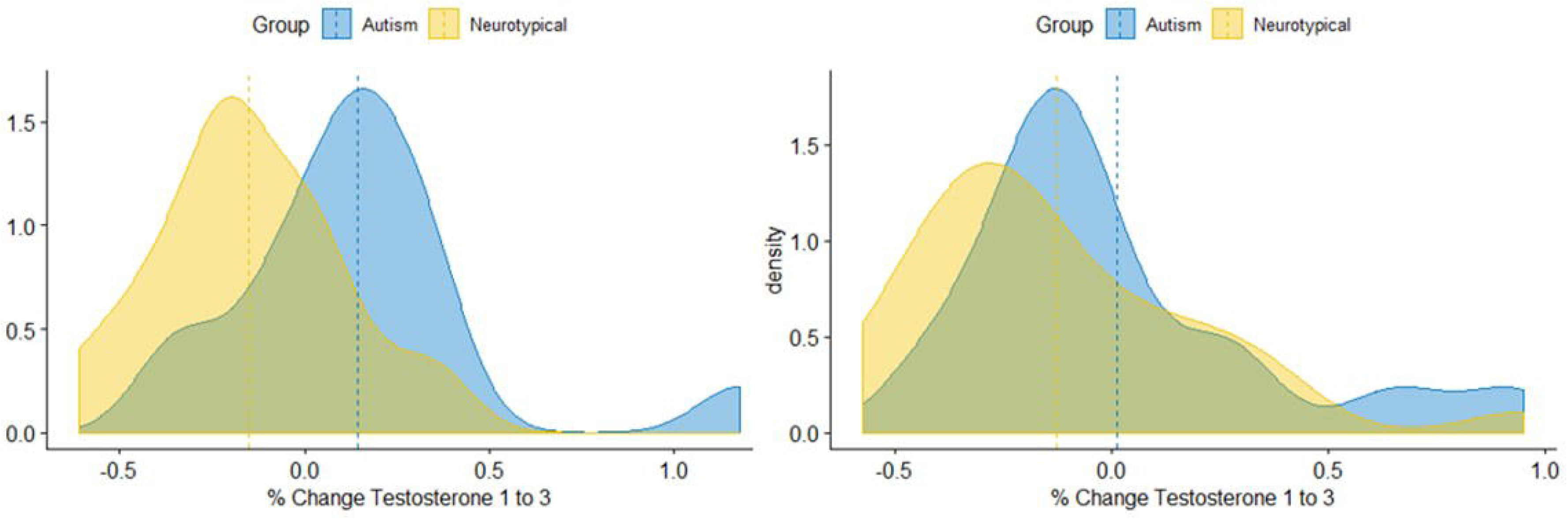
Density plots of %change testosterone from time 1 to 3 under the oxytocin (left panel) and placebo condition (right panel) between groups (Autism and Typical). The dashed lines indicate the means for each group.

Robust regression was performed to ensure that the overall results were not driven by hormone level changes for a small number of highly influential individuals. For oestradiol %change from time point 1 to 3, 8 of 90 observations were substantially down-weighted (< 0.60, Huber weight), which did not change the effect of group in the model. For testosterone %change from time point 1 to 3, 5 of 90 observations were substantially down-weighted (< 0.60, Huber weight), which again did not change the overall results. Further highlighting the distinct patterns of testosterone change under the oxytocin condition, 11 of 16 autistic women (69%) showed a testosterone increase, whereas only 8 of 29 (28%, including 3 with an increase < 5%) typical women showed a testosterone increase.

## Discussion

In the present study, we sought to identify interactions between oxytocin and steroid hormones that could potentially influence the outcomes of oxytocin administration in experimental and clinical settings. Given the underrepresentation of women in both autism research and endocrinology research, this work has made several important contributions to a sparse literature. First, on average, women in our study showed small but significant decreases in both salivary oestradiol and testosterone levels over time, which is consistent with expected diurnal rhythms of sex steroid hormone [40]. However, post-hoc tests revealed that decreases in hormone levels over time were limited to typical women, with autistic women instead showing average increases. Oxytocin enhanced the between-group difference in %change testosterone, with the majority of autistic women (11 of 16) showing an increase and the majority of typical women (21 of 29) showing a decrease over time. The pattern for autistic women is in line with findings by Gossen et al. [8] that oxytocin increased men’s testosterone levels, which peaked 120 minutes after administration. In a study by Weisman et al. [7] on the effects of oxytocin administration on parent-child interactions, fathers’ testosterone levels decreased over time, but those who received oxytocin had significantly higher testosterone levels at 40, 60, and 80 minutes after administration relative to the placebo condition. The pattern observed in autistic women is consistent with the idea of a masculinized phenotype [32,33], as they showed sex hormone responses to oxytocin that are more like those of typical males. In the typical women in our study, we see no evidence that oxytocin elevated testosterone levels relative to placebo— rather, testosterone levels were slightly higher under the placebo condition (~5 min post-administration: 56.4±18.9 Placebo, 52.7±17.3 Oxytocin; ~90 min post-administration: 54.7±15.6 Placebo, 52.4±16.2 Oxytocin).

As testosterone levels tended to increase under both the placebo and oxytocin conditions in autistic women, this begs the question of why. Endogenous testosterone levels are reported to increase in contexts related to competition and mating [41], neither of which is applicable to our experimental context. One possibility is that autistic women experienced more stress during fMRI scanning, as elevated testosterone levels have been reported in response to social and physical stress [42]. How oxytocin—which is generally considered to have anxiolytic effects [43]—may influence stress hormone levels in humans is presently unknown. However, if oxytocin deficiencies do exist in autism, oxytocin may enhance stress response, as shown in a rodent model of oxytocin-deficient female mice [44]. Grillon et al. [45] demonstrated that oxytocin can increase anxiety in humans, specifically defensive response to an unpredictable threat. Therefore, it is possible that the testosterone increase observed in our autistic group was related to heightened stress response. If oxytocin indeed has anxiogenic effects in certain individuals, they will be unlikely to benefit from therapeutic interventions involving oxytocin.

Furthermore, our data presented a rare opportunity to compare salivary hormone levels between autistic and typical adult women. Ruta et al. [25] and Xu et al. [46] reported no difference in oestradiol levels in blood between a mixed sex population of autistic adults (33 men, 25 women) vs. controls and 61 mothers of autistic children vs. control mothers, respectively. Ruta et al. [25] further reported no difference in testosterone or the testosterone to oestradiol ratio between autistic and typical individuals. By contrast, Schwarz et al. [47] reported elevated testosterone levels in 23 women with Asperger’s syndrome relative to controls, and Xu et al. found elevated testosterone levels in mothers of autistic children relative to control mothers [46]. While our study has the advantage of comparing testosterone levels in two samples collected at least one week apart, which should be a more reliable baseline than a single measure as used in the studies described above, our study was underpowered to detect a small difference in testosterone levels between autistic and typical women due to the small sample size and high variability in salivary testosterone.

Recent findings that medical conditions associated with elevated testosterone, such as polycystic ovary syndrome, are more common in autistic women [24,48] supports potential roles of dysregulation of sex steroid hormone systems in autism, even if such differences are not reliably found in blood or saliva samples. Although it did not meet the threshold for statistical significance, the difference in testosterone levels between the autistic and typical women in our study was in the expected direction (see Table 1).

Interestingly, the ratio of testosterone to oestradiol was positively correlated with autistic-like traits in our overall study sample, and negatively correlated with self-reported empathy. Although the correlations are modest, they are broadly consistent with previous reports of associations between hormone measures and social traits in children and women. Prenatal testosterone levels, as measured in amniotic fluid, are positively and negatively correlated, respectively, with scores on the childhood versions of the Autism-Spectrum Quotient [49] and Empathy Quotient [50]. Adult women with elevated testosterone levels due to congenital adrenal hyperplasia are reported to have higher levels of autistic-like traits than controls [51], whereas girls with lower testosterone levels scored higher on cognitive empathy than girls with higher testosterone [52]. While differences in prenatal sex steroid hormones have been posited to contribute to sex differences in autistic-like traits and empathy [53], these data suggest that postnatal hormone levels contribute to individual differences in psychological traits within one sex and that consideration of multiple hormones within one pathway may be a better representation of this relationship.

This study has several limitations. First, as this was primarily a neuroimaging experiment, participants were scheduled based on scanner availability rather than within a narrow time window typically used for hormone studies. However, time of day may not significantly affect salivary testosterone levels in women [54] and, if it does, the expectation is a decrease across the day. Participants were scheduled during the luteal phase of the menstrual cycle, and we relied on self-report for this determination. Similarly, use of hormonal contraceptive was self-reported, although these variables may not strongly influence testosterone levels in women [54]. Lastly, given the relatively small sample size and lack of previous studies assessing post-oxytocin administration hormone changes in women, further studies are needed to confirm whether typical and autistic women show opposite patterns of sex steroid hormone changes in response to oxytocin, with autistic women showing a pattern of increase in testosterone more like that previously reported in typical men.

Notably, not all autistic women showed increased testosterone levels and not all typical women showed decreased testosterone levels after oxytocin administration. Whether individual changes in endogenous hormone levels are related to the effects of oxytocin on behavior—particularly the social behaviours that oxytocin administration aims to enhance in clinical populations—is yet to be tested. In a recent oxytocin trial in autistic children, Parker et al. [55] found that low pre-treatment baseline oxytocin levels were predictive of improvement in social functioning with oxytocin. In an oxytocin administration study of typical women, oxytocin administration was found to decrease response time to face stimuli, but only among women with high endogenous testosterone levels [10]. Given the inconsistent findings of oxytocin clinical trials to date [56,57] and the seemingly opposite roles of oxytocin and testosterone in neuropsychiatric conditions including autism and schizophrenia [11], the possibility that baseline sex steroids or oxytocin-associated changes in sex steroids could serve be a biomarker of response may help to identify autistic people expected to benefit most from interventions involving oxytocin.

## Acknowledgements

Salivary testosterone and oestradiol assays were performed by the NIHR Cambridge Biomedical Research Centre, Core Biochemical Assay Laboratory

## References

1. Johnson Z V., Young LJ. Oxytocin and vasopressin neural networks: Implications for social behavioral diversity and translational neuroscience. Neurosci Biobehav Rev. 2017.

2. Meyer-Lindenberg A, Domes G, Kirsch P, Heinrichs M. Oxytocin and vasopressin in the human brain: social neuropeptides for translational medicine. Nat Rev Neurosci. 2011;12:524–538.

3. Bakermans-Kranenburg MJ, van I Jzendoorn MH. Sniffing around oxytocin: review and meta-analyses of trials in healthy and clinical groups with implications for pharmacotherapy. Transl Psychiatry. 2013;3:e258.

4. Quintana DS, Smerud KT, Andreassen OA, Djupesland PG. Evidence for intranasal oxytocin delivery to the brain: Recent advances and future perspectives. Ther Deliv. 2018. 2018. https://doi.org/10.4155/tde-2018-0002.

5. Bethlehem RAI, Baron-Cohen S, van Honk J, Auyeung B, Bos PA. The oxytocin paradox. Front Behav Neurosci. 2014;8:48.

6. Weisman O, Feldman R. Oxytocin administration affects the production of multiple hormones. Psychoneuroendocrinology. 2013;38:626–627.

7. Weisman O, Zagoory-Sharon O, Feldman R. Oxytocin administration, salivary testosterone, and father-infant social behavior. Prog Neuro-Psychopharmacology Biol Psychiatry. 2014;49:47–52.

8. Gossen a, Hahn a, Westphal L, Prinz S, Schultz RT, Gründer G, et al. Oxytocin plasma concentrations after single intranasal oxytocin administration - a study in healthy men. Neuropeptides. 2012;46:211–215.

9. Weisman O, Schneiderman I, Zagoory-Sharon O, Feldman R. Salivary vasopressin increases following intranasal oxytocin administration. Peptides. 2013;40:99–103.

10. Holtfrerich SKC, Schwarz KA, Sprenger C, Reimers L, Diekhof EK. Endogenous testosterone and exogenous oxytocin modulate attentional processing of infant faces. PLoS One. 2016;11:1–19.

11. Crespi BJ. Oxytocin, testosterone, and human social cognition. Biol Rev. 2016;91:390–408.

12. Insel TR, Young L, Witt DM, Crews D. Gonadal steroids have paradoxical effects on brain oxytocin receptors. J Neuroendocrinol. 1993;5:619–628.

13. Okabe S, Kitano K, Nagasawa M, Mogi K, Kikusui T. Testosterone inhibits facilitating effects of parenting experience on parental behavior and the oxytocin neural system in mice. Physiol Behav. 2013;118:159–164.

14. Frayne J, Nicholson HD. Effect of Oxytocin on Testosterone Production by Isolated Rat Leydig Cells is Mediated Via a Specific Oxytocin Receptor1. Biol Reprod. 1995. 1995. https://doi.org/10.1095/biolreprod52.6.1268.

15. Jirikowski GF, Caldwell JD, Pilgrim C, Stumpf WE, Pedersen CA. Changes in immunostaining for oxytocin in the forebrain of the female rat during late pregnancy, parturition and early lactation. Cell Tissue Res. 1989. 1989. https://doi.org/10.1007/BF00218899.

16. McCarthy MM, McDonald CH, Brooks PJ, Goldman D. An anxiolytic action of oxytocin is enhanced by estrogen in the mouse. Physiol Behav. 1996;60:1209–1215.

17. Dai D, Li QC, Zhu Q Bin, Hu SH, Balesar R, Swaab D, et al. Direct involvement of androgen receptor in oxytocin gene expression: Possible relevance for mood disorders. Neuropsychopharmacology. 2017;42:2064–2071.

18. Dumais KM, Veenema AH. Vasopressin and oxytocin receptor systems in the brain: Sex differences and sex-specific regulation of social behavior. Front Neuroendocrinol. 2016;40:1–23.

19. APA. American Psychiatric Association, 2013. Diagnostic and statistical manual of mental disorders (5th ed.). 2013.

20. Parker KJ, Garner JP, Libove R a., Hyde S a., Hornbeak KB, Carson DS, et al. Plasma oxytocin concentrations and OXTR polymorphisms predict social impairments in children with and without autism spectrum disorder. Proc Natl Acad Sci. 2014:1402236111-.

21. Feldman R, Golan O, Hirschler-Guttenberg Y, Ostfeld-Etzion S, Zagoory-Sharon O. Parent-child interaction and oxytocin production in pre-schoolers with autism spectrum disorder. Br J Psychiatry. 2014;205:107–112.

22. Bakker-Huvenaars MJ, Greven CU, Herpers P, Wiegers E, Jansen A, van der Steen R, et al. Saliva oxytocin, cortisol, and testosterone levels in adolescent boys with autism spectrum disorder, oppositional defiant disorder/conduct disorder and typically developing individuals. Eur Neuropsychopharmacol. 2018.

23. Baron-Cohen S, Auyeung B, Nørgaard-Pedersen B, Hougaard DM, Abdallah MW, Melgaard L, et al. Elevated fetal steroidogenic activity in autism. Mol Psychiatry. 2015. 2015. https://doi.org/10.1038/mp.2014.48.

24. Cherskov A, Pohl A, Allison C, Zhang H, Payne RA, Baron-Cohen S. Polycystic ovary syndrome and autism: A test of the prenatal sex steroid theory. Transl Psychiatry. 2018. 2018. https://doi.org/10.1038/s41398-018-0186-7.

25. Ruta L, Ingudomnukul E, Taylor K, Chakrabarti B, Baron-Cohen S. Increased serum androstenedione in adults with autism spectrum conditions. Psychoneuroendocrinology. 2011. 2011. https://doi.org/10.1016/j.psyneuen.2011.02.007.

26. Phaik Ooi Y, Weng SJ, Kossowsky J, Gerger H, Sung M. Oxytocin and Autism Spectrum Disorders: A Systematic Review and Meta-Analysis of Randomized Controlled Trials. Pharmacopsychiatry. 2017.

27. Winslow J, Insel T. Social status in pairs of male squirrel monkeys determines the behavioral response to central oxytocin administration. J Neurosci. 1991;11(7):2032–2038.

28. Rilling JK, Demarco AC, Hackett PD, Chen X, Gautam P, Stair S, et al. Sex differences in the neural and behavioral response to intranasal oxytocin and vasopressin during human social interaction. Psychoneuroendocrinology. 2014;39:237–248.

29. Gao S, Becker B, Luo L, Geng Y, Zhao W, Yin Y, et al. Oxytocin, the peptide that bonds the sexes also divides them. Proc Natl Acad Sci. 2016;113:7650–7654.

30. Baron-Cohen S, Wheelwright S, Skinner R, Martin J, Clubley E. The Autism Spectrum Quotient□: Evidence from Asperger syndrome/high functioning autism, males and females, scientists and mathematicians. J Autism Devl Disord. 2001;31:5–17.

31. Baron-Cohen S, Wheelwright S. The Empathy Quotient: An Investigation of Adults with Asperger Syndrome or High Functioning Autism, and and Normal Sex Differences.pdf. J Autism Dev Disord. 2004. 2004.

32. Baron-Cohen S, Lombardo M V., Auyeung B, Ashwin E, Chakrabarti B, Knickmeyer R. Why are Autism Spectrum conditions more prevalent in Males? PLoS Biol. 2011;9.

33. Greenberg DM, Warrier V, Allison C, Baron-Cohen S. Testing the Empathizing-Systemizing theory of sex differences and the Extreme Male Brain theory of autism in half a million people. Proc Natl Acad Sci U S A. 2018. 2018. https://doi.org/10.1073/pnas.1811032115.

34. Bethlehem RAI, Lombardo M V., Lai MC, Auyeung B, Crockford SK, Deakin J, et al. Intranasal oxytocin enhances intrinsic corticostriatal functional connectivity in women. Transl Psychiatry. 2017;7:e1099–8.

35. Wakabayashi A, Baron-Cohen S, Wheelwright S, Goldenfeld N, Delaney J, Fine D, et al. Development of short forms of the Empathy Quotient (EQ-Short) and the Systemizing Quotient (SQ-Short). Pers Individ Dif. 2006;41:929–940.

36. Gröschl M. Current status of salivary hormone analysis. Clin Chem. 2008;54:1759–1769.

37. Sollberger S, Ehlert U. How to use and interpret hormone ratios. Psychoneuroendocrinology. 2016. 2016. https://doi.org/10.1016/j.psyneuen.2015.09.031.

38. Fox, J. & Weisberg S. Robust Regression in R: An Appendix to An R Companion to Applied Regression. 2nd edition. Sage. 2011. 2011.

39. Team RDC, R Development Core Team R. R: A Language and Environment for Statistical Computing. R Found Stat Comput. 2016. 2016. https://doi.org/10.1007/978-3-540-74686-7.

40. Dabbs JM. Salivary testosterone measurements: Reliability across hours, days, and weeks. Physiol Behav. 1990;48:83–86.

41. Zilioli S, Bird BM. Functional significance of men’s testosterone reactivity to social stimuli. Front Neuroendocrinol. 2017.

42. Taylor SE, Klein LC, Lewis BP, Gruenewald TL, Gurung RAR, Updegraff JA. Biobehavioral responses to stress in females: Tend-and-befriend, not fight-or-flight. Psychol Rev. 2000. 2000. https://doi.org/10.1037/0033-295X.107.3.411.

43. MacDonald K, Feifel D. Oxytocin’s role in anxiety: A critical appraisal. Brain Res. 2014;1580:22–56.

44. Amico JA, Mantella RC, Vollmer RR, Li X. Anxiety and stress responses in female oxytocin deficient mice. J Neuroendocrinol. 2004. 2004. https://doi.org/10.1111/j.0953-8194.2004.01161.x.

45. Grillon C, Krimsky M, Charney DR, Vytal K, Ernst M, Cornwell B. Oxytocin increases anxiety to unpredictable threat. Mol Psychiatry. 2013.

46. Xu XJ, Shou XJ, Li J, Jia MX, Zhang JS, Guo Y, et al. Mothers of Autistic Children: Lower Plasma Levels of Oxytocin and Arg-Vasopressin and a Higher Level of Testosterone. PLoS One. 2013;8.

47. Schwarz E, Guest PC, Rahmoune H, Wang L, Levin Y, Ingudomnukul E, et al. Sex-specific serum biomarker patterns in adults with Asperger’s syndrome. Mol Psychiatry. 2011. 2011. https://doi.org/10.1038/mp.2010.102.

48. Pohl A, Cassidy S, Auyeung B, Baron-Cohen S. Uncovering steroidopathy in women with autism: A latent class analysis. Mol Autism. 2014. 2014. https://doi.org/10.1186/2040-2392-5-27.

49. Auyeung B, Baron-Cohen S, Ashwin E, Knickmeyer R, Taylor K, Hackett G. Fetal testosterone and autistic traits. Br J Psychol. 2009. 2009. https://doi.org/10.1348/000712608X311731.

50. Chapman E, Baron-Cohen S, Auyeung B, Knickmeyer R, Taylor K, Hackett G. Fetal testosterone and empathy: evidence from the empathy quotient (EQ) and the ‘reading the mind in the eyes’ test. Soc Neurosci. 2006. 2006. https://doi.org/10.1080/17470910600992239.

51. Knickmeyer R, Baron-Cohen S, Fane BA, Wheelwright S, Mathews GA, Conway GS, et al. Androgens and autistic traits: A study of individuals with congenital adrenal hyperplasia. Horm Behav. 2006. 2006. https://doi.org/10.1016/j.yhbeh.2006.02.006.

52. Pascual-Sagastizabal E, Azurmendi A, Sánchez-Martín JR, Braza F, Carreras MR, Muñoz JM, et al. Empathy, estradiol and androgen levels in 9-year-old children. Pers Individ Dif. 2013. 2013. https://doi.org/10.1016/j.paid.2013.01.019.

53. Baron-Cohen Simon S. Empathizing, systemizing, and the extreme male brain theory of autism. vol. 186. 2010.

54. Liening SH, Stanton SJ, Saini EK, Schultheiss OC. Salivary testosterone, cortisol, and progesterone: Two-week stability, interhormone correlations, and effects of time of day, menstrual cycle, and oral contraceptive use on steroid hormone levels. Physiol Behav. 2010. 2010. https://doi.org/10.1016/j.physbeh.2009.10.001.

55. Parker KJ, Oztan O, Libove RA, Sumiyoshi RD, Jackson LP, Karhson DS, et al. Intranasal oxytocin treatment for social deficits and biomarkers of response in children with autism. Proc Natl Acad Sci. 2017:201705521.

56. Guastella AJ, Hickie IB. Oxytocin Treatment, Circuitry, and Autism: A Critical Review of the Literature Placing Oxytocin into the Autism Context. Biol Psychiatry. 2016.

57. Keech B, Crowe S, Hocking DR. Intranasal oxytocin, social cognition and neurodevelopmental disorders: A meta-analysis. Psychoneuroendocrinology. 2018;87:9–19.

