## Supplementary Tables and Figures for "Salivary sex hormone levels following oxytocin administration in autistic and typical women"

**Table S1**. Comparison of demographic characteristics, psychological questionnaire scores, and baseline hormone levels in the Autism and Typical groups excluding participants who reported taking hormonal contraceptives (n=8). Values are mean ± SD, unless otherwise specified.

|  | **Autism** | **Typical** | ***p-value*** |
| --- | --- | --- | --- |
| **n** | 16 | 21 |  |
| **Demographics** |  |  |  |
| **Age (years)** | 29.9 ± 8.4 | 28.7 ± 9.0 | 0.55 |
| **Full-IQ^1^** | 126.4 ± 20.0 | 112.3 ± 14.7 | 0.12 |
| **Psychological variables** |  |  |  |
| **Autism-Spectrum Quotient (AQ)** | 37.1 ± 5.1 | 13.8 ± 8.1 | < 0.01** |
| **Empathy Quotient (EQ)** | 20.7 ± 11.7 | 57.7 ± 12.6 | < 0.01** |
| **Baseline hormone levels^2^** |  |  |  |
| **Baseline oestradiol (pg/ml)** | 1.0 ± 0.3 | 1.2 ± 0.5 | 0.19 |
| **Baseline testosterone (pg/ml)** | 70.3 ± 24.9 | 69.4 ± 21.4 | 0.91 |

^1^ Wechsler Abbreviated Scale of Intelligence
^2^ Baseline hormone levels were calculated as the mean of the two pre-administration samples per participant.
** p < 0.01

**Table S2**. ANOVA of Group × Drug condition effect on salivary oestradiol levels from time point 1 (before administration) to time point 2 (~5 min post-administration).

^1^ Type II sum of squares.

|  | **Sum Sq^1^** | **df** | **F** | **p** |
| --- | --- | --- | --- | --- |
| **Drug condition (Oxytocin or Placebo)** | 0.0005 | 1 | 0.0061 | 0.9378 |
| **Group (Autism or Typical )** | 0.1541 | 1 | 1.9113 | 0.1705 |
| **Drug condition × Group** | 0.0464 | 1 | 0.5754 | 0.4502 |
| **Residual** | 6.7726 | 84 |  |  |

**Table S3**. ANOVA of Group × Drug condition effect on salivary testosterone levels from time point 1 (before administration) to time point 2 (~5 min post-administration).

|  | **Sum Sq^1^** | **df** | **F** | **p** |
| --- | --- | --- | --- | --- |
| **Drug condition (Oxytocin or Placebo)** | 0.0006 | 1 | 0.0121 | 0.9125 |
| **Group (Autism or Typical)** | 0.0882 | 1 | 1.685 | 0.1977 |
| **Drug condition × Group** | 0.093 | 1 | 1.7247 | 0.1926 |
| **Residual** | 4.5018 | 86 |  |  |

^1^ Type II sum of squares.

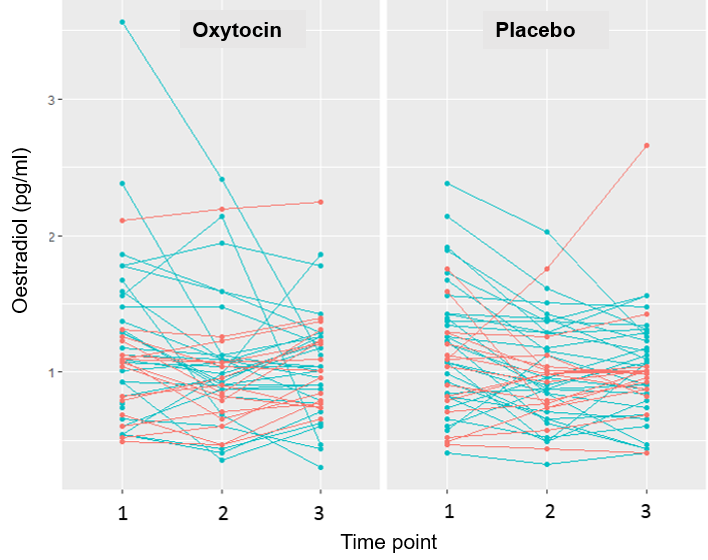

**Figure S1.** Salivary oestradiol levels across the three time points (1 = baseline, 2 = ~5 min post-administration, 3 = ~90 min post-administration) each participant under oxytocin (left) and placebo (right) drug conditions. Autistic participants (Autism group) are indicated in red while typical participants (Typical group) are in blue.

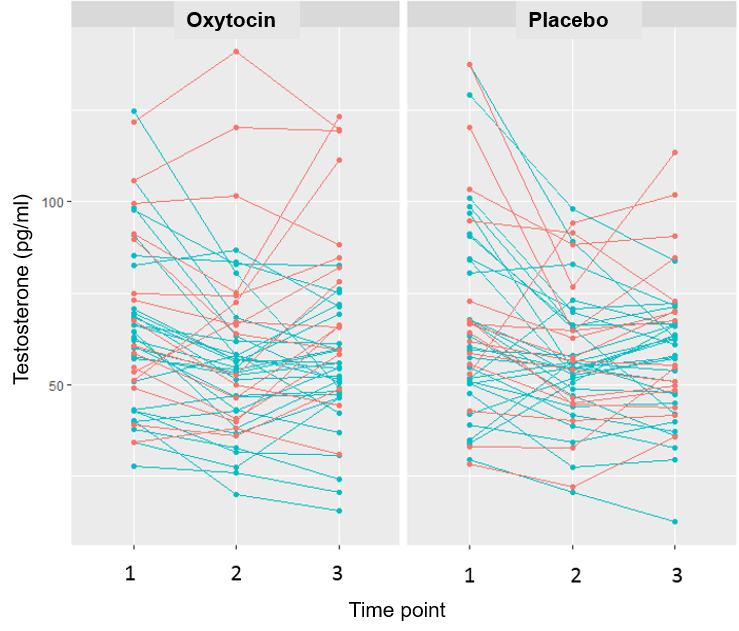

**Figure S2.** Salivary testosterone levels across the three time points (1 = baseline, 2 = ~5 min post-administration, 3 = ~90 min post-administration) each participant under oxytocin (left) and placebo (right) drug conditions. Autistic participants (Autism group) are indicated in red while typical participants (Typical group) are in blue.
